# Real-time chromatin dynamics at the single gene level during transcription activation

**DOI:** 10.1101/111179

**Authors:** Thomas Germier, Silvia Kocanova, Nike Walther, Aurélien Bancaud, Haitham Ahmed Shaban, Hafida Sellou, Antonio Zaccaria Politi, Jan Ellenberg, Franck Gallardo, Kerstin Bystricky

## Abstract

Genome dynamics relate to regulation of gene expression, the most fundamental process in biology. Yet we still do not know whether the very process of transcription drives spatial organization and chromatin conformation at specific gene loci. To address this issue, we have optimized the ANCHOR/ParB DNA labeling system for real-time imaging and quantitative analysis of the dynamics of a single-copy transgene in human cells. Transcription of the transgene under the control of the endogenous Cyclin D1 promoter was induced by addition of 17β-estradiol. Motion of the ANCHOR3-tagged DNA locus was recorded in the same cell prior to and during appearance of nascent mRNA visualized using the MS2 system. We found that transcription initiation resulted in rapid confinement of the mRNA-producing gene. The confinement was maintained even upon inhibition of pol2 elongation. It did not occur when recruitment of pol2 or transcription initiation was blocked by anti-estrogens or Triptolide. These results suggest that preinitiation complex formation and concomitant reorganization of the chromatin domain constrains freedom of movement of an induced gene’s promoter within minutes. Confined diffusion reflects assembly of functional protein hubs and DNA processing during the rate-limiting steps of transcription.

## Introduction

3D organization of the genome contributes significantly to regulation of major nuclear processes. Changes in average position of chromosome loci in a population of cells correlate with local or global changes in DNA metabolism (Therizols et al. 2014;Taddei et al. 2006;Kocanova et al. 2010;Cabal et al. 2006;Chambeyron and Bickmore 2004;Osborne et al. 2004;Schuettengruber and Cavalli 2009;Robinett et al. 1996;Chuang et al. 2006a). This is notably the case for gene transcription, where active genes tend to associate with clusters of RNA polymerase II (pol2) (Feuerborn and Cook 2015). By imaging pol2, its cofactors and mRNA these transcription hubs have been shown to be relatively immobile (Kimura et al. 2002;Ghamari et al. 2013;Cisse et al. 2013;Darzacq et al. 2007), but the motion of the associated DNA has not been reported. Consequently, neither do we know if the observed reduced protein mobility is an intrinsic property of the transcription machinery or an indirect effect of changes in chromatin conformation, nor what the precise kinetics of this reorganization at short timescales are.

Indeed, real time analysis of chromatin at short time scales relevant for the analysis of transcription activation (minutes) has been hampered by methodological limitations. Existing technologies to visualize DNA loci usually rely on highly repetitive sequences, based on insertion of hundreds of repeats of bacterial operator sequences to which fluorescent repressor fusion proteins bind with high affinity (called FROS for fluorescent repressor operator system (Straight et al. 1996)), or using multiplexed short guide RNAs that stably recruit catalytically inactive dCas9-GFP fusion proteins to a large, repetitive genomic region and partially unwind the target DNA sequence (Chen et al. 2013;Ma et al. 2015). These technologies confirmed that transcription impacts nuclear localization of gene domains. However, they do not allow tagging of genes within immediate vicinity of regulatory elements by fear to disturb their very function. Nevertheless, it was shown that, in yeast, the mobility of a gene was increased by permanently recruiting the potent activator VP16 or chromatin remodeling factors (Neumann et al. 2012). This effect could stem from constitutive local decondensation of chromatin near the labelled gene. In mouse ES cells, in contrast, it was reported that in the presence of trans-activation by expressing Nanog, overall gene motion was reduced (Ochiai et al. 2015). In both studies gene motion was compared in different cells. To truly assess immediate changes in chromatin motion during transcription activation, DNA dynamics of a single-copy gene has to be analyzed in real-time while simultaneously monitoring steps of mRNA synthesis in the same cell.

To achieve this, we developed a novel ANCHOR (ParB/INT) DNA labeling system (ANCHOR3) for use in human cells. Stable insertion of the ANCHOR3 system into cells in which transcription of target genes can be activated under physiological conditions enables fluorescence imaging of a single locus in the same cell over time, without interfering with gene expression.

We demonstrate that the optimized ANCHOR3 system in combination with the MS2 system is ideally suited for simultaneous visualization of DNA and mRNA at a single gene level in living human cells at high spatio-temporal resolution. We show that transcription initiation, not elongation, constrains local displacement of the hormone-induced Cyclin D1 gene as an immediate response to the transcription process in human cells.

## Results

To simultaneously visualize DNA and mRNA of a gene, we labeled a Cyclin D1 (CCND1) transgene with a new, improved ANCHOR3 system (see materials and methods). The ANCHOR system was derived from prokaryotic chromosome partitioning components and originally implemented in yeast (Saad et al. 2014). Specific association of a few ParB/OR protein dimers to a limited number of parS binding sites within the bacterial chromosome′s partitioning site initiates formation of a large nucleoprotein complex dependent on non-specific, dynamic ParB/OR binding and ensuing oligomerization (Passot et al. 2012;Graham et al. 2014;Sanchez et al. 2015). The ANCHOR system thus relies simply on a short ANCH/INT sequence (<1kb) that can be inserted immediately adjacent, within a few base-pairs, to regulatory elements.

The transgene is further composed of the endogenous CCND1 promoter, a CCND1 cDNA cassette including its 3′enhancer region and 24 repeats of the MS2 MCP protein-binding sequence within the CCND1 3′-UTR (Yunger et al. 2010) (Fig. 1a). The construct was inserted into an FRT site within the genome of estrogen receptor *alpha* (ER*α*)-positive MCF-7 human mammary tumor cells (Fig. 1a). In the engineered, monoclonal cells, called ANCH3-CCND1-MS2, fluorescent OR3-fusion proteins form a single focus at the ANCH3 site of the transgene that can be readily tracked in real time (Fig. 1b; Videos S1, S2). To characterize the binding kinetics of OR3 proteins at and around the ANCH3 site, we used fluorescence recovery after photo-bleaching (FRAP) of OR3-EGFP labeled spots (Fig. 1b). Association and dissociation of OR3-EGFP at the ANCH3-tagged site was in a dynamic steady state with a measured half-life of 57±2s (Fig. 1b). To estimate the copy number of OR3 proteins at an ANCH3 site in steady state, we performed confocal imaging calibrated by fluorescence correlation spectroscopy (FCS), which allowed us to convert pixel fluorescence intensity to protein concentration or number (Fig. S1a-e). The fluorescence intensity of the ANCHOR3 spot increased with OR3-EGFP abundance (Fig. S1f). Under our OR3 expression conditions, we calculated an average of 481 ± 274 fluorescent molecules per site, corresponding to a significant amplification of the nine OR3 dimers bound specifically to the parS sites of the ANCH3 sequence (Fig. S1f).

**Figure 1:**
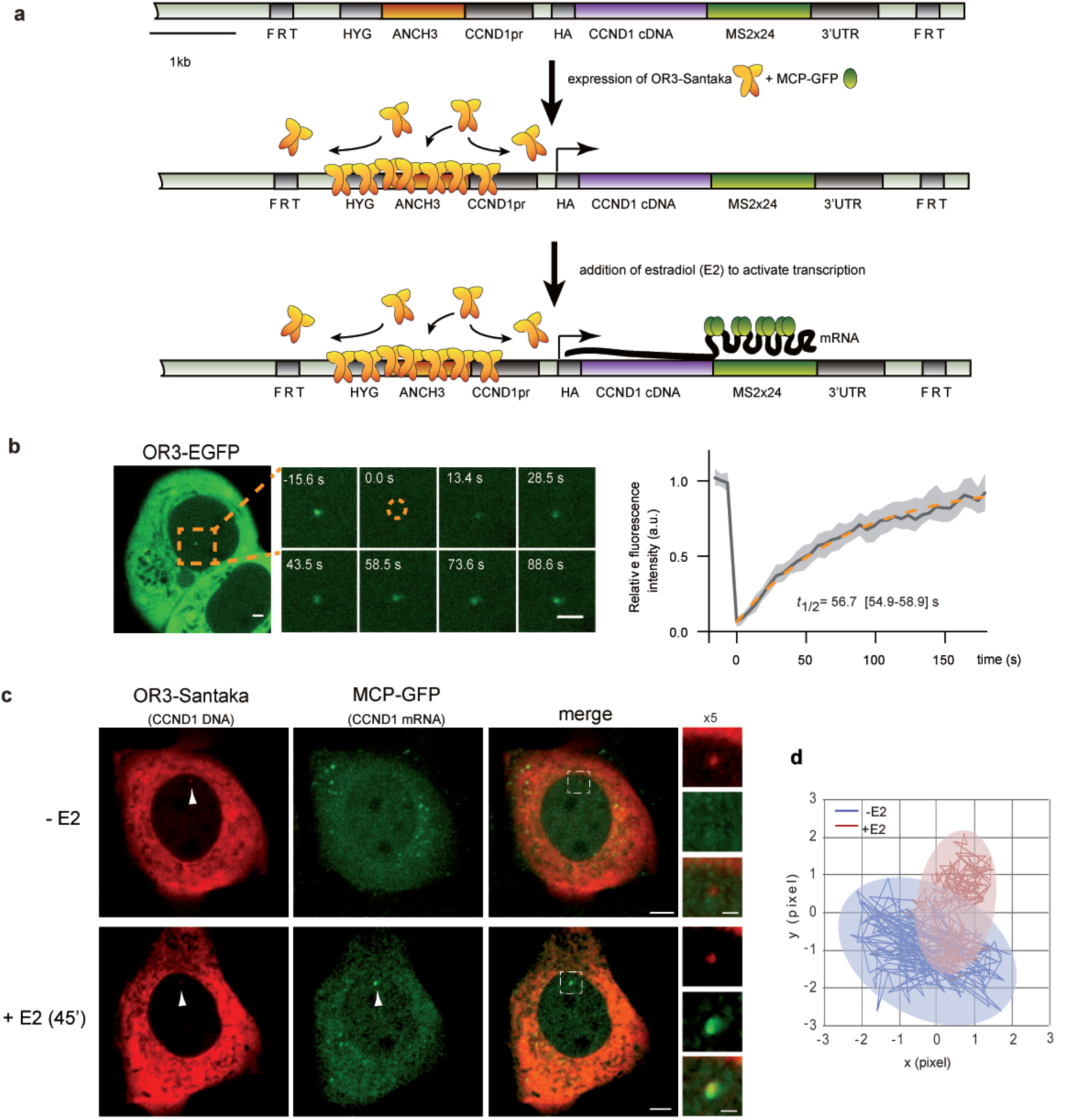
Real-time visualization of a single Cyclin D1 (CCND1) gene locus in human cells. (a) Schematic representation of a stably inserted construct (ANCH3-CCND1-MS2) comprising the CCND1 gene under its endogenous promoter, adjacent to a unique ANCH3 sequence, 24x MS2 repeats within the 3’UTR and a hygromycin selection gene (HYG). The construct is flanked by FRT sites for integration into MCF-7 FRT cells. Transient transfection with OR3 and MCP-tagged fluorescent proteins results in their accumulation at the ANCH3 and MS2 sequences (after estradiol (E2) stimulation), respectively (raw 3D images in supplemental video 1).(b) Fluorescent spots are easily detectable in transfected cells. A representative cell with an OR3-EGFP spot is shown. Region imaged during fluorescence recovery after photobleaching (FRAP) is indicated in orange. At time t=0s a circular region enclosing the ANCHOR spot was bleached and fluorescence recovery of the spot was followed over time. Relative fluorescence intensity (RFI) was calculated according to (see Materials and methods online and Fig. S1; right panel; solid line: mean, shadowed region: lower and upper quartile; n = 44 cells, 4 experiments with n ≥ 6 cells per experiment). Data were fitted to a single exponential. The 95% confidence interval is indicated in brackets. Scale bar: 2 μm. (c) Representative images of transiently transfected ANCH3-CCND1-MS2 cells expressing OR3-Santaka and MCP-EGFP (raw images in videos S1, S2). CCND1 DNA (red spot) colocalizes with transcribed mRNA (green spot) as MCP-EGFP associates with MS2 stem loops 45min after adding 100nM estradiol (E2). The same cell is shown before and after addition of E2. Scale bar = 5μm and 2μm (for cropped images), (d) Example of 2D trajectories and area explored over 50s (250ms acquisition, 200 steps) of the OR3-Santaka labeled CCND1 locus recorded before (-E2) and after (+E2) transcription activation.

The MCP-EGFP signal corresponding to accumulated CCND1 transcripts is detectable near the ANCHOR3-labeled DNA site 45 minutes after 17β-estradiol (E2) addition to G1-synchronized cells grown in steroid-stripped media (Fig. 1c; video S2), consistent with the fact that ERα target gene expression is triggered 10 minutes after E2 addition (Hah et al. 2011;Métivier et al. 2003). Thus, the engineered cell line enabled real-time imaging of a single copy gene during transcription activation by the endogenous ERα, under physiological conditions.

To directly test whether changes in gene expression impact local chromatin dynamics, we recorded the motion of the fluorescent ANCHOR3-Santaka-tagged gene in the same cell prior to and 45 min after adding 100nM E2, while monitoring the appearance of MCP-EGFP-labelled mRNA signals (Fig. 1c). Live cell tracking revealed that movement of the CCND1 gene is locally constrained upon induction of its transcription (Fig. 1d, table 1). We quantified the average displacement of the tagged transgene by plotting the mean square displacement (MSD) to each time interval ∆t (see materials and methods). MSD curves calculated from time-lapse image series acquired with an inter-frame interval of 250 ms for a total of 50 s are shown in Fig. 2a. We found that MSD plots of E2-activated CCND1 differed significantly from those of non-activated cells (Fig.2a left panels, Fig. S2, S3). In contrast, E2 had no effect on the behavior of a non-genic ANCH3-only construct integrated at the same genomic location (Fig. 2a right panels, Fig. S2, S3), confirming that the measured decline in mobility was due to transcription of the transgene rather than to unspecific, genome-wide effects of hormone addition. We also examined the motion of the ANCH3-CCND1-MS2 or ANCH3-only constructs inserted into distinct chromosomes (G7, A11 and D11 clones; Fig. 2a, Fig. S2). At the single cell level, we recorded large variations in MSD of the transgene in all clones, but these variations did not correlate with any specific insertion site (Fig. S2). Intriguingly, motion of the ANCH3-tagged single gene locus followed two distinct regimes with an increase in the slope of the recorded MSD at time intervals >5 s in these human mammary tumor cells synchronized in G1 (Fig. 2b left panel, Fig. S2). The average MSD curves over 21 trajectories followed a non-linear, anomalous diffusive behavior characteristic of objects moving in complex environments such as the nucleoplasm (Saxton 2009)(Fig. S2). Hence, we analyzed these MSDs on the basis of a generalized diffusion model obeying a power law *MSD~kv^α^* with *k* as prefactor, *t* the time interval, and *α* the anomalous diffusion exponent, a model shown to reflect chromatin motion in several cell types and under various conditions (Manzo and Garcia-Parajo 2015). Applying this model to our data, we demonstrate that at short <5 s time intervals diffusion of the chromatin fiber of a single gene domain is highly anomalous (*α* <0.4) and subjected to local constraints. At greater time intervals, the slope of the population averaged MSD curve increases (*α*~1) suggesting that the fiber of the non-transcribed gene locus (ANCH3-CCND1-MS2 or ANCH3 only) is rather mobile, almost freely diffusing. In contrast, the dynamic behavior of the mRNA producing CCND1 loci differs significantly: the MSD slope remains low (a ~0.5) consistent with significant confinement of the actively transcribed locus (Fig. 2b).

**Figure 2:**
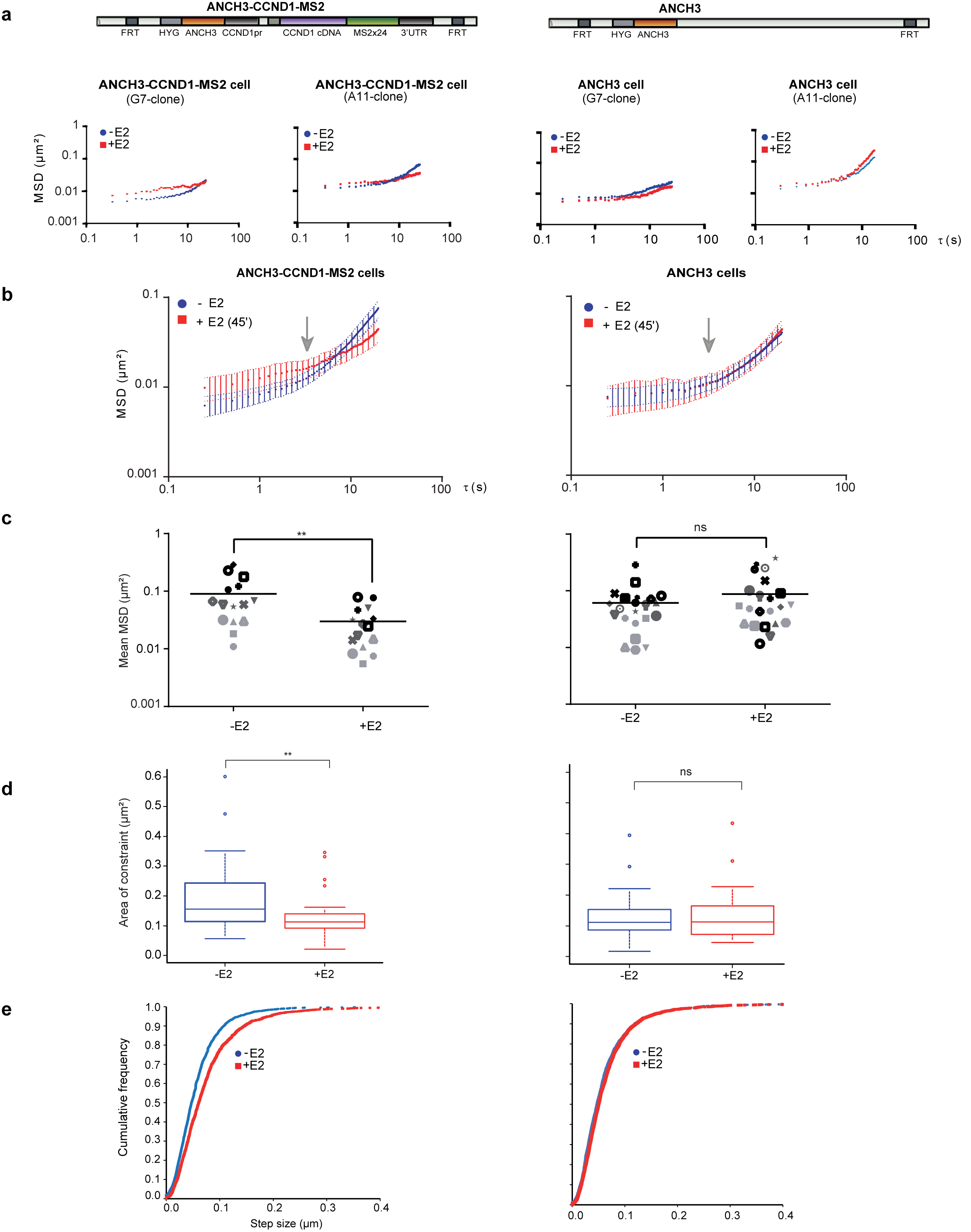
Cyclin D1 motion is confined during estradiol induced transcription activation. Representative single cell MSD curves of OR3-Santaka loci in ANCH3-CCND1-MS2 and ANCH3 cells at two different chromosomal insertion sites (clones G7 and A11; distinct stable FRT insertions in MCF-7 cells) before and 45 minutes after E2 addition. (b) Averaged MSD curves of OR3-Santaka loci tracking in ANCH3-CCND1-MS2 (n=14) and ANCH3 (n=14) cells before and after E2 addition. The arrow indicates the 5 s time point separating two diffusive regimes (Fig. S4 b). (c) Average squared displacement between 2 s and 40 s (mean MSD) of tracking of the OR3-Santaka spot in ANCH3-CCND1-MS2 cells (left panel, n=15, p-value=0.006) and ANCH3 cells (right panel, n=21, p-value=0.3) before and 45 min after E2 addition. (d) Area of constraint of the OR3-Santaka locus in ANCH3-CCND1-MS2 cells (left panel, n=24, p-value=0.006) and ANCH3 cells (right panel, n=20, p-value=0.860) before and after E2 addition. The boxes show the median and 25–75 percentiles of the data. Asterisks indicate data points beyond the 95th percentile. Student’s t-test: *P-values: >0.05 (ns), <0.05 (*), <0.01 (**), <0.001 (***), <0.0001 (****). (e) Cumulative distribution function of CCND1 loci step size in ANCH3-CCND1-MS2 cells (left panel, n=14, p-value<0.001) and ANCH3 cells (right panel, n=14, p-value=0.03) before and after E2 addition.

**Table 1:** Summary of confined areas and speeds calculated for different treatments. Data are shown as average +/-standard deviation. The first line contains all cells that were not treated (0min, n=63) or treated 45 min with E2 (45min, n=34), independently of the rest of experiment.

| | Treatment | Area of confinement ($\mu\text{m}^2$) | | | | | Speed ($\mu\text{m/s}$) | | | | |
| --- | --- | --- | --- | --- | --- | --- | --- | --- | --- | --- | --- |
|  |  | 0 min | 30 min | 45 min | 75 min | 105 min | 0 min | 30 min | 45 min | 75 min | 105 min |
| ANCH3-CCND1-MS2 cells | all E2 | 0.175 ± 0.119<br>N=63 | | 0.117 ± 0.06<br>N=34 (E2) | | | 0.255 ± 0.085<br>N=63 | | 0.293 ± 0.131<br>N=34 (E2) | | |
| | E2 (45') | 0.195 ± 0.134<br>N=24 | | 0.133 ± 0.082<br>(E2) | | | 0.231<br>N=14 | | 0.304<br>(E2) | | |
| | E2 (105') | 0.106 ± 0.05<br>N=4 | | 0.106 ± 0.06<br>(E2) | 0.129 ± 0.06<br>(E2) | 0.101 ± 0.05<br>(E2) | 0.260 | | 0.263<br>(E2) | 0.261<br>(E2) | 0.323<br>(E2) |
| | (E2+DRB) | 0.143 ± 0.054<br>N=8 | | 0.121 ± 0.041<br>(E2) | 0.172 ± 0.150<br>(E2+DRB) | 0.173 ± 0.106<br>(E2+DRB) | 0.215 | | 0.246<br>(E2) | 0.259<br>(E2+DRB) | 0.318<br>(E2+DRB) |
| | (DRB+E2) | 0.163 ± 0.075<br>N=7 | 0.254 ± 0.114<br>(DRB) | | 0.205 ± 0.24<br>(DRB+E2) | | 0.276 | 0.270<br>(DRB) | | 0.305<br>(DRB+E2) | |
| | E2 then DRB | 0.155 ± 0.033<br>N=3 | | 0.123 ± 0.039<br>(E2) | 0.124 ± 0.036<br>(DRB) | 0.163 ± 0.040<br>(DRB) | 0.227 | | 0.269<br>(E2) | 0.251<br>(DRB) | 0.308<br>(DRB) |
| | DRB (105') | 0.148 ± 0.040<br>N=5 | | 0.190 ± 0.064<br>(DRB) | 0.160 ± 0.058<br>(DRB) | 0.109 ± 0.061<br>(DRB) | 0.262 | | 0.273<br>(DRB) | 0.341<br>(DRB) | 0.306<br>(DRB) |
| | (E2+TPL) | 0.154 ± 0.057<br>N=3 | | 0.099 ± 0.045<br>(E2) | 0.213 ± 0.055<br>(E2+TPL) | 0.168 ± 0.016<br>(E2+TPL) | 0.321 | | 0.417<br>(E2) | 0.330<br>(E2+TPL) | 0.333<br>(E2+TPL) |
| | (TPL+E2) | 0.219 ± 0.189<br>N=10 | 0.143 ± 0.083<br>(TPL) | | 0.253 ± 0.179<br>(TPL+E2) | | 0.296 | 0.288<br>(TPL) | | 0.321<br>(TPL+E2) | |
| | no treatment | 0.118 ± 0.035<br>N=6 | | 0.112 ± 0.066 | 0.126 ± 0.068 | 0.085 ± 0.053 | 0.253 | | 0.251 | 0.271 | 0.236 |
| ANCH3 cells | E2 (45') | 0.132 ± 0.090<br>N=20 | | 0.137 ± 0.096<br>(E2) | | | 0.242<br>N=14 | | 0.256<br>(E2) | | |

To fully exploit our ability to track a single gene, we computed an average squared displacement from 2s to 40 s (mean MSD) before and after induction of transcription in the same cell. For both constructs, in the absence of E2, the mean MSD ranged from 0.009 to 0.290 μm^2^ (Fig. 2c). This heterogeneity in the amplitude of motion is coherent with variations in the nuclear environment owing to crowding in mammalian nuclei (Hancock 2004;Huet et al. 2014;Ochiai et al. 2015). Independently of the initial mobility, mean MSDs recorded for each cell producing MCP-labeled mRNA were consistently reduced after addition of E2 to ANCH3-CCND1-MS2 cells (Fig.2c left panel; between 0.006 and 0.090μm^2^; n=15; p-value=0.006). In contrast, variations in mean MSD of the non-genic ANCH3 locus were not significantly affected by E2 addition (Fig. 2c right panel; n=21; p-value=0.3).

To more accurately describe the behavior of the tracked chromatin locus, we determined its area of confinement, track length and speed in addition to time averaged MSDs which suffer from approximating experimental errors (Kepten et al. 2015). We found that spatial confinement of the transcribed locus reflected obstructed diffusion. The nuclear area explored by the tagged locus over a 50 s time interval in the absence of hormone was reduced by 33% upon addition of E2, from 0.175 ±0.119 μm^2^ (n=63, no E2, no detectable MCP labeled mRNA) to 0.117 ±0.006μm^2^ (n=34; visible mRNA accumulation at the tagged locus after 45 min E2) (Fig. 2d, table 1). Surprisingly, the mean step size of the tracked locus was greater after E2 addition (Fig. 2e) and, as a consequence, the apparent velocity of the transcribed CCND1 locus increased from 0.26 μm/s to 0.29 μm/s (table 1). The ANCH3-only locus did not alter its speed in the presence of E2 (table 1).

Confinement of the chromatin fiber could stem from steric hindrance due to protein loading or a change in the physical parameters of the fiber, or both (Banks and Fradin 2005). We first tested whether recruitment of large transcriptional co-factor complexes by hormone-bound ERα to the CCND1 promoter could influence chromatin mobility. Similarly to E2, Tamoxifen (OH-Tam) triggers ERα binding to responsive promoters, which leads to recruitment of numerous proteins and chromatin remodeling complexes; in contrast to E2, OH-Tam-bound ERα attracts transcriptional co-repressors (Shang et al. 2000;Liu and Bagchi 2004). However, changes in the recorded amplitude of the MSD calculated from tracking the ANCH3-tagged CCND1 locus after addition of 1μM OH-Tam did not show a distinct trend. Motion was variable and similar to the dynamic behavior of the constructs in the absence of hormone (Fig. S3). We conclude that association of a multitude of transcriptional co-factors recruited by OH-Tam-bound ERa alone, in the absence of transcription, cannot explain local confinement of the chromatin fiber.

We next assessed the role of pol2 activity on motion of the tagged CCND1 locus using two distinct pol2 inhibitors (Fig. 3). E2-stimulated cells were treated with an elongation-inhibitor, the adenosine analogue 5,6-dichloro-1-β-D-ribofuranosylbenzimidazole (DRB). CCND1 mRNA signals disappeared 30 min after addition of 50μM DRB to ANCH3-CCND1-MS2 cells indicating that completion of elongation, i.e. transcription of the 3’UTR including MS2 repeats, was efficiently inhibited (Fig. 3b). The appearance of the MSD plot of the CCND1 locus tracking after DRB addition to cells maintained in E2-containing medium was similar to the one in cells which had not been treated with DRB (Fig. 3b, single cell MSD curves), suggesting that confined motion was sustained despite blocking elongation. Mean MSD values and area of confinement for several cells analyzed 45 min after E2 stimulation and 30 min after adding DRB did also not change significantly (Fig. 3c, table 1). When pre-treating cells for 30 min by DRB prior to adding hormone, we again observed a rapid decline in CCND1 motion upon mRNA production in every single cell analyzed (Fig. 3d). It is further known that in DRB-treated cells, transcription initiation by pol2 is preserved but elongation aborts rapidly within the first transcribed exon (Gribnau et al. 1998). Our observations let us speculate that initiating but not elongating pol2 confines chromatin dynamics locally. To confirm this hypothesis, we analyzed the tagged gene’s motion in cells treated with 500 nM Triptolide (TPL), an inhibitor of TFIIH that blocks pol2 at the promoter after PIC assembly (Vispé et al. 2009;Jonkers et al. 2014) (Fig. 3a). Addition of TPL to E2-stimulated mRNA-producing cells released the constraint as the MCP-GFP signal diffuses away from the ANCH3-tagged transgene (Fig. 3e). Indeed, mean MSD values of the ANCHOR3 spot’s track increased after 30 min or 60 min depending on efficiency of TPL for evicting pol2 from the gene body (Fig. 3f, table 1). Addition of E2 to TPL pre-treated cells had no impact on the recorded motion of the transgene (Fig. 3g, table 1). Hence, our data suggest that events linked to PIC initiation induce changes in chromatin mobility, leading to increased local velocity at time scales > 5s within a largely confined area.

**Figure 3:**
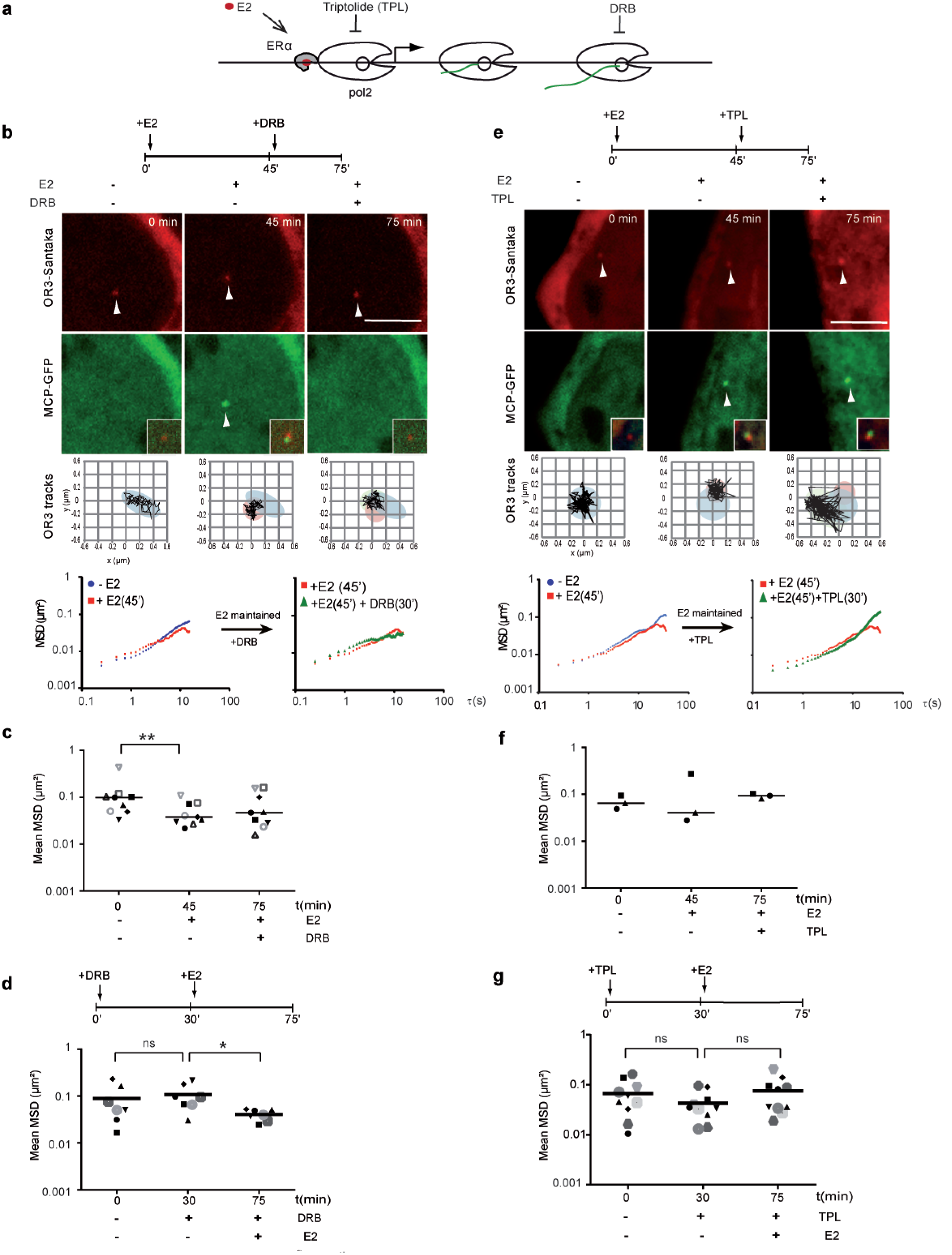
Transcription initiation but not elongation confines chromatin motion. (a) Schematic representation of the steps of the E2-induced ERa-bound CCND1 the inhibited by Triptolide and DRB during transcription initiation or elongation, respectively, (b, e) Experimental set-up, 100 nM of E2 was used to activate transcription and 50 pM DRB to block elongation (b) or 500 nM of Triptolide (TPL) to block initiation of transcription (e). The CCND1 locus was tracked at time points before and after treatment as specified. Representative images from live cell imaging of ANCH3-CCND1-MS2 cells (clone G7) treated first with E2 for 45 min and then with DRB or TPL for 30 min, respectively (b, e). mRNA production is activated by E2 and then blocked by inhibiting transcription elongation as evidenced by the appearance and disappearance of the MCP-EGFP signal. Scale bar = 5 pm. Successive positions of the OR3-Santaka spot are displayed in the bottom squares (122 steps, 250 ms interval). Representative single cell MSD curves of OR3 loci detected in a cell treated with E2 and then DRB. (c) Average squared displacement between 2 s and 40 s (mean MSD) of single cells (n=9) treated successively with E2 and then DRB (d) Mean MSD (250 ms interval. 200 steps, n=7) of single cells pre-treated with 50 pM DRB for 30 min (p-value=0.578) and then treated with 100 nM E2 for 45 min (p-value=0.031). (e, f) as (b, c), but after timed treatment with TPL. (g) Mean MSD (250 ms interval, 200 steps, n=10) of single cells pre-treated with 500 nM TPL for 30 min (p-value=0.117) and then treated with 100 nM E2 for 45min (p-value=0.065).

## Discussion

Real-time tracking of a single-copy Cyclin D1 gene in the same cell prior to and during hormone-induced mRNA synthesis revealed that transcription initiation rapidly confines the mRNA-producing gene and alters its diffusive behavior. Confinement was maintained even upon inhibition of pol2 elongation, but did not occur when recruitment of pol2 or transcription initiation was blocked by anti-estrogens or Triptolide. Our results suggest that PIC formation and concomitant reorganization of the chromatin domain constrain freedom of movement of an induced gene′s promoter within minutes, compatible with the establishment of a transcriptional hub. Indeed, pol2 aggregates in numerous rather immobile foci (Cisse et al. 2013;Darzacq et al. 2009). Several models concur in saying that a small fraction of active pol2 forms clusters with reduced mobility as transcription initiates (Stasevich et al. 2014;Kimura et al. 2002;Cisse et al. 2013). This clustering is dependent on the presence of initiating pol2 complexes (Mitchell and Fraser 2008). But what causes pol2 to stop moving? In principle, our observation that transcription initiation locally confines chromatin dynamics within minutes is compatible with the idea that pol2 foci assemble at active genes, and that, as the transcription initiation bubble forms, a decline in DNA freedom of movement leads to reduced pol2 mobility.

Regulation of the CCND1 locus has been shown to involve intragenic looping (Dalvai et al. 2013). Such conformational changes in gene domain organization, similar to those observed during glucocorticoid stimulated transcription of MMTV tandem array gene loci in mouse adenocarcinoma cells (Stavreva et al. 2015), are likely to have direct consequences for chromatin dynamics. For instance, the anchoring of several chromosome fibers within pol2 foci may increase the drag coefficient and hence reduce chromatin displacements. The changes in local dynamics we describe are thus compatible with reorganization of pre-existing chromatin folding within the gene domain at the 100 kb range via long-range looping (Mourad et al. 2014;Chuang et al. 2006b;Therizols et al. 2014). Greater local velocity of the transcribed locus might increase the frequency of interaction with transcriptional cofactors and polymerases of the gene within its regulatory compartment similar to what was recently modelled as a ‘nanoreactor’(Haddad et al. 2017). Increased collisions are compatible with the formation of gene domain specific chromatin clustering (also called ‘topologically associated domains’ or TADs) readily detected by crosslinking methods in mammary tumor cells (unpublished;(Le Dily et al. 2014;Giorgetti et al. 2014). In turn, confined dynamics may prevent formation of unwanted long range contacts as transcription proceeds.

At the sub-megabase level, TADs comprise one or a few open reading frames and their regulatory elements (gene domains) (Ulianov et al. 2016), particularly in human mammary tumor cells (Barutcu et al. 2016;Mourad et al. 2014;Le Dily et al. 2014;Fullwood et al. 2009)(Kocanova et al. in preparation). If the existence of TADs is elusive in yeast, increased ligation frequencies also occur around gene bodies at the 2 kb range (Hsieh et al. 2015). Most of our knowledge of chromatin dynamics stems from work in budding yeast (Bystricky 2015;Botstein and Fink 2011;Wang et al. 2015;Taddei and Gasser 2012). In particular, live cell chromatin motion of a series of tagged genomic yeast loci fits a Rouse model of polymer dynamics, in which the MSD increases with time with a power-law scaling and an anomaly exponent α~0.5 (Hajjoul et al. 2013). Assuming that within a ~100 kb chromatin domain around any of the tagged sites, at least one gene is actively transcribed in a population of yeast nuclei, the reported dynamic behavior characterizes active chromatin. In agreement, obstructed diffusion characterizes the active CyclinD1 transgene locus in human cells here. In the absence of transcription the tagged single human transgene domain was highly dynamic, nearly freely diffusing. Similarly, our unpublished observations in yeast evidence increased motion when mutating RNA pol 2 (Mathon, Wang et al. in preparation). Transcription-induced, pol2 dependent, intra-domain contacts therefore likely result from the apparent highly diffusive behavior of chromatin in living cells. Anomalous diffusion was also reported for telomeres (Bronshtein et al. 2015) and for gene arrays (Annibale and Gratton 2015) but changes were derived from comparing motion in cells monitored under different conditions. The computed MSD curves of these loci characterized by specific structures differ from the ones we compute for a single gene locus which emphasizes the need for future modeling to better define physical parameters of the chromatin fiber in human cells.

Because dynamic properties of chromatin have been implicated in all fundamental cellular processes we propose that decrease in chromatin motion as a consequence of transcription initiation confers essential functions to chromatin dynamics. The powerful real-time imaging approach of a single DNA locus undergoing functional changes presented here using the ANCHOR system is widely applicable to other loci and genomes for studying rapid biological processes with single cell resolution, and completing the picture emerging from imaging RNA pol2 and mRNA, but also from chromosome conformation capture data.

## Materials and Methods

### Cell line

The human breast cancer cell line MCF-7 (purchased from ATCC) was used to generate stable FRT/LacZeo clones. Cells were grown in Dulbecco’s modified Eagle medium F-12 (red DMEM/F12) completed with 10% FBS (Gibco), 1% Sodium Pyruvate (Gibco) and 0.5% Gentamycine (Gibco) or in phenol red free DMEM/F12 completed with 10% charcoal stripped serum, 1% Sodium Pyruvate (Gibco) and 0.5% Gentamycine (Gibco), in a water-saturated atmosphere containing 5% CO2 at 37°C. Transfections were carried out using FuGENE HD Transfection Reagent (Promega), according to the supplier’s recommendation.

### ANCHOR3 labeling system

ANCH3 corresponds to a specific chromosome partition sequence and was amplified directly by PCR from the genome of an undisclosed exotic bacteria *(upon acceptance, name will be disclosed and vectors made available for purchase from Addgene).* The PCR product was then cloned using BamH1/HindIII into pCDNA FRT vector digested by BglU/HindIM (Invitrogen). Insertion was verified by ApaI digestion and sequencing. OR3 corresponding to the cognate ParB protein was amplified by PCR and cloned via BglII/KpnI directly into pGFP-c1 digested by the same enzyme. Insertion was verified by digestion and sequencing. Functionality of the construct was verified by co-transfection of both vectors in Hela cells, and produced clearly identifiable fluorescent spots. To construct the ANCH3-CCND1-MS2 transgene (Fig. 1a) the ANCH3 sequence was amplified by PCR with primers containing EcoRV restriction sites and ligated into the EcoRV digested pCDNA5-FRT/CCND1pr-HA-CCND1/24MS2/3′UTR plasmid (kindly provided by YaronShav-Tal (Yunger et al. 2010)).

### Engineering of stable FRT/LacZeo clones expressing the CCND1 transgene

MCF-7 cells were transfected using pFRT//lacZeo kit from Invitrogen according to the supplier′s protocol. Selection of FRT clones was performed using 75 μg/ml Zeocin (InvivoGen) in red DMEM/F12 medium. Selection medium was renewed every 3 days. Three clones, showing the same growing behavior and ERα-target gene expression profiles as MCF-7 cells were kept for use: G7, A11 and D11. Clones G7, A11, D11 were transfected with 1 μg of ANCH3-CCND1-MS2 or ANCH3 plasmids and 6 μg of plasmid pOG44 encoding Flipase (Invitrogen). Selection of positive clones was performed with 75 μg/ml Hygromycin (Invitrogen) and presence of a single fluorescent focus was verified by fluorescence microscopy 24h after cell transfection with 500ng OR3-Santaka. ANCH3-CCND1-MS2 cells as well as control cells with only an ANCH3 insertion were maintained in red DMEM/F12 completed with 75 μg/ml Hygromycin.

### Fluorescence correlation spectroscopy (FCS) and fluorescence recovery after photobleaching (FRAP) experiments

For FCS and FRAP experiments, 2 × 10^4^cells were seeded on 8well Lab-TekI chambered cover glass (Nunc). To visualize spots, cells were transfected 1h after cell seeding with a plasmid encoding OR3-EGFP using FuGENE HD transfection reagent (Promega) according to the manufacturer’s instructions for a 3:1 transfection reagent:DNA ratio. To express free mEGFP as a control for FCS, cells were transfected 1h after cell seeding with a plasmid encoding mEGFP (kindly provided by J. Lippincott-Schwartz) using FuGene6 transfection reagent (Promega) according to the manufacturer’s instructions for a 3:1 transfection reagent:DNA ratio. FCS and FRAP experiments were performed 24 to 48h after cell seeding/transfection in CO_2_-independent imaging medium (Gibco, custom-made) supplemented with 20% (v/v) FBS, 1% (v/v) sodium pyruvate and 1% (v/v) L-glutamine.

### Fluorescence recovery after photobleaching (FRAP)

Imaging was performed on a Zeiss LSM780 ConfoCor3 confocal microscope using a 40x, 1.2 NA, water Korr FCS objective. Prior to each FRAP time-lapse, an overview image of the measured interphase cell with a nuclear spot was taken. GFP was excited with a 488 nm laser (Ar, 25 mW, 0.4% AOTF transmission, 2.3 μW at probe) and detected with a GaAsP detector using a 492-552 nm detection window (Ax, Ay = 83 nm, ∆*z* = 0.4, 60×60 pixels, 9 z-slices). 40 images were acquired at a 5 s time interval. After three prebleach images a circular ROI (12 pixels wide) containing the ANCHOR spot was bleached with five iterations and maximal laser power. For bleaching controls, cells with spots were acquired with the same conditions as in the FRAP experiments but without bleaching the fluorescent spot. The z-position was stabilized using Zeiss Definite Focus. Due to hardware limitations the first four images have a time interval between 3.4 s and 6.6 s. These reproducible differences were taken into account in the analysis. The image analysis was performed using FiJi (http://fiji.sc/Fiji). For each FRAP time-course the position of the OR3-EGFP ANCHOR spot was tracked in 3D using the FiJi MOSAIC plugin for 3D single-particle tracking (http://mosaic.mpi-cbg.de/?q=downloads/imageJ,(Sbalzarini and Koumoutsakos 2005)). If a spot was not detectable after photobleaching, its position was interpolated. The mean fluorescence intensity *F_s_* was measured in a 5×5 rectangular region around the tracked spot. The background fluorescence intensity *F_bg_*was measured from a ROI in the unbleached part of the nucleus. Relative fluorescence intensity *RI* at time point *t_i_* (Fig. S1d) is given by

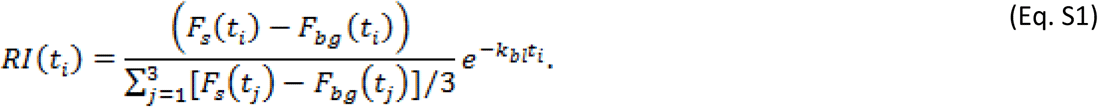

whereby the data was normalized to the average of the first three pre-bleach images and bleach corrected using an exponential factor. The bleaching rate *k_bl_* was estimated from the FRAP time-courses without bleaching the ANCHOR3 spots. Overall bleaching was below 10% within the acquired 40 frames. The post-bleach kinetics were fitted to a single exponential function as described in the legend to Fig. S1d.

### Fluorescence correlation spectroscopy (FCS) and protein number estimation

Imaging and photon counts were acquired on a Zeiss LSM780 ConfoCor3 confocal microscope using a 40×, 1.2 NA, water Korr FCS objective. Cells were imaged (∆*x*, ∆*y* = 83 nm, ∆*z* = 0.4, 164×164 pixels, 9 z-slices) and subsequently two positions in the nucleus outside of ANCHOR3 spots were selected for FCS at the z-position of the central slice. For imaging, GFP was excited with a 488 nm laser (Ar, 25 mW, 0.4% AOTF transmission, 2.3 μW at probe) and detected with a GaAsP detector using a 492-552 nm detection window. For pre-FCS imaging the same detector settings, pixel size, pixel dwell time, and laser settings were used as for FRAP, allowing the conversion of fluorescence intensities in the first pre-bleach FRAP images to concentrations and protein numbers based on an FCS calibration (Eq. S3). For FCS, GFP was excited with a 488 nm laser (Ar, 25 mW, 0.05% AOTF transmission, 0.28μW at probe) and the photon counts were recorded for 30 s using an avalanche photodiode detector (APD; 505-590 nm detection). To estimate the effective volume, a water solution of Alexa488 (Lifetech) with a known diffusion coefficient (*D*_alexa_ = 441 μm^2^sec^−1^, M. Wachsmuth, EMBL, personal communication) was measured before each experiment.

The raw photon counts were processed using FluctuationAnalyzer 4G (http://www.embl.de/~wachsmut/downloads.html). This program computes the autocorrelation function (ACF), correction factors, e.g. due to background, and fits the ACF to physical models of diffusion (see Wachsmuth et al., 2015, for further details). The ACF *G*(*τ*) was fit to

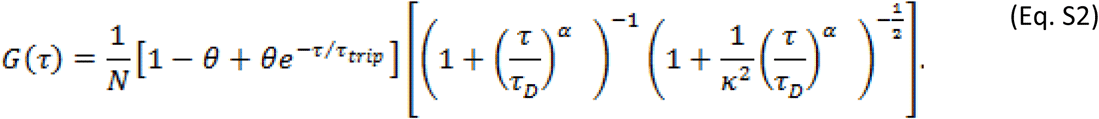

Equation S2 describes anomalous diffusion: *N* denotes the number of particles in the effective volume, *κ=* 5.5 the structure factor, i.e. the ratio of axial to lateral radius of the effective volume, *τ*_D_the characteristic diffusion correlation time, α the anomaly parameter. The parameter *θ* is the fraction of molecules in a non-fluorescent state and *τ_trip_* the apparent life-time in this state. The value of *τ_trip_* was set to 100 μs and the other parameters were fit to the data.

The concentration of molecules in the effective focal volume *V_eff_* is given by *C = N/V_eff_N_A_* where 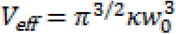 and *N_A_* is the Avogadro′s constant. The lateral focus radius *w*_0_ is given by 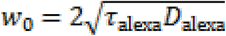 where *τ_alexa_* is the characteristic diffusion correlation time measured for Alexa488. The counts per molecule are given by *CPM* = *N/〈I〉* where *〈I〉* is the average photon counts.

For each experiment, a calibration curve (Fig. S1c) was calculated from the mean fluorescence intensity (FI) and the concentration obtained from FCS. The mean FI was measured in a 5×5 px large square at the location of the FCS measurement. The data points were fit to a line that describes the relationship between fluorescence intensity *FI* and concentration *C*:

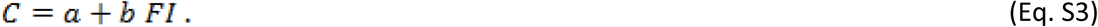

Based on Eq. S3, each pixel of images acquired with the same settings as the pre-FCS images can be converted to a concentration. The protein number in each voxel was obtained by multiplying the concentration *C* with the voxel volume *V_vox_* and *N_A_* (see Fig. S1e).

To estimate the protein number at the ANCHOR site, the first pre-bleach image of the FRAP time-course was used. The FI was converted to protein number per pixel using the calibration curve in Eq. S3 and *V_vox_.*

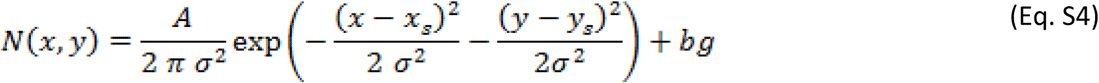

In a 14×14 rectangular region at the location of the ANCHOR3 site a 2D Gaussian was fit (Eq. S4), where *x_s_* and *y_s_* are the coordinates of the spot, *bg* the background nuclear signal and *A* the total protein number at the Anchor site. For the fit the Gauss Fit on Spot ImageJ plugin (http://imagej.nih.gov/ij7plugins/gauss-fit-spot/index.html) was used.

### Fluorescence live cell imaging under stimulation

For live cell tracking during transcription activation, 80 000 cells were plated in 35mm glabsosttom culture plates (Ibidi, Biovalley) in red DMEM/F12 medium and allowed to attach for 24h. Red DMEM/F12 medium was changed for phenol red free DMEM/F12 medium supplemented with 10% charcoal stripped serum and cells were kept in the latter medium for 72h. Cells were co-transfected with 500ng OR3-Santaka and 1μg MCP-GFP DNA vectors 24h before observation.

To activate transcription of the ANCH3-CCND1-MS2 transgene, cells were maintained in L-15 medium (Liebovitz’s, Gibco) supplemented with 10% charcoal stripped serum, a buffer appropriate for live cell imaging.

Observations were made using a Nipkodwisk confocal system (Revolution; Andor) installed on a microscope (IX-81), featuring a confocal spinning disk unit (CSU22;Yokogawa) and a cooled electron multiplying chargecoupled device camera (DU 888; Andor). The system was controlled using the Revolution IQ software (Andor). Images were acquired using a 60× Plan Apo 1.42 oil immersion objective and a two-fold lens in the optical path. Single laser lines used for excitation werep udmiopded solid-state lasers exciting GFP fluorescence at 488 nm (50 mW; Coherent) and Santaka fluorescence at 561 nm, and a Quad band pass emission filter (Di0T1405/488/56 8/647-13×15×0,5; Semrock) allowed collection of the green and redfluorescence. Pixel size was 110 nm. Movies containing 200 image frames acquired with an exposure time of 250ms were recorded. Images were processed using ICY and FIJI software.

After imaging of the cells without transcription stimulation, 100nM 17β-estradiol (E2; Sigma) was added directly under the microscope and dynamics of the OR3-Santaka spot in the same cells were recorded 45min later. Before acquisition under E2 stimulation, presence of the MCP-GFP foci, indicating active transcription elongation, was verified in each analyzed cell (Fig. 1c). To block transcription elongation, we added fresh L-15 medium containing 50μM DRB (5,6-dichloro-1-β-D-ribofuranosylbenzimidazole; Sigma). The cells were imaged 30min after DRB addition and disappearance of the MCP-GFP foci was verified (Fig. 3b). To examine the impact of transcription initiation on chromatin motion in living cells, we added fresh L-15 medium containing 500nM Triptolide (TPL, Sigma) before or after stimulation by E2. The cells were imaged under the same conditions as for DRB treatment. To study the effect of OH-Tamoxifen (OH-Tam; Sigma), cells were maintained in L-15 medium and imaged before and 45min after 1μM OH-Tam treatment.

### Lateral drift test

Lateral drift was analyzed by cross correlation. Cross correlation assumes that shape of the structures imaged in the sample is not expected to change significantly during the acquisition; the structure itself can be used to determine whether spatial shift between subsequent images exists. For single cell analysis, movies were cropped and registered by the ImageJ plug-in StackReg using translation and rigid body functions for drift correction, before tracking of the DNA locus using ICY tracker (2). The translation function calculates the amount of translation (A**r**) by a vectoring analysis from **x** = **r** + ∆**r**. Where, **x** and **r** are the output and input coordinates. Rigid body transformation is appropriate because coordinates are **x** = {{cos *ϑ*, −sin *ϑ*}, {sin *ϑ*, cos *ϑ*}} · **r** + ∆**r**, considering both the amount of translation (∆**r**) and the rotation by an angle *ϑ* (2).

### Mean square displacement

Particle tracking experiments and MSD calculations were carried out using ICY and MatLab software. Tracks of 200 frames were scored. OR3-Santaka spots were detected and tracked using the Spot detector and spot tracking plug-in from ICY in single cells. Mean square displacements (MSD) were calculated in Matlab using the following equation:

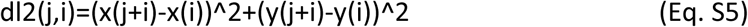

Averaged MSD resulted from averaging the MSD at each time interval (Fig. 2b, n=14). Mean MSD were extracted from the average squared displacement from 2s to 40 s of the ANCH3 locus during the time of acquisition in one cell (Fig. 2c, 3c, 3d, 3f and 3g, n=15,n=21, n=9, n=7, n=3 and n=10 respectively). Areas of confinement were obtained using the raw trajectories over 50sec and fitted based on an ellipse.

### Statistics

Results were analyzed using two different tests. A Student’s t-test with a confidence interval of 95% was used for data presented in Fig. 2c, 2d and 2e. A Wilcoxon signed-rank test was used for data presented in Fig. 3c, 3d, 3f and 3g

## Acknowledgements

We thank Y.Shav-Tal for plasmid pcDNA5/FRT/GOI-MS2. M.Dalvai is acknowledged for help during the initial phases of the study. We acknowledge support from the TRI-Imaging platform Toulouse. TG was supported by a doctoral fellowship from the Ligue Nationale Contre le Cancer. KB was supported by HFSPO RGP0044, ANR ANDY, Fondation ARC. JE was supported by EMBL and the EU-FP7-SystemsMicroscopy NoE (258068) and the 4DN NIH Common Fund (U01 EB021223), NW by the EMBL International PhD Programme (EIPP).

## Author contributions

Performed experiments: TG, SK, FM

Analysed data: TG, SK, AB, HSh, KB

NW performed and analysed, AZP analysed and JE supervised the FRAP and FCS analysis.

Generated material: TG (cell lines), HSe (cell lines, constructs), FG (ANCHOR3 system)

Wrote paper: KB, TG, SK;

## References

Annibale P, Gratton E. 2015. Single cell visualization of transcription kinetics variance of highly mobile identical genes using 3D nanoimaging. Sci Rep 5: 9258. http://www.nature.com/doifinder/10.1038/srep09258.

Banks DS, Fradin C. 2005. Anomalous Diffusion of Proteins Due to Molecular Crowding. Biophys J 89: 2960–2971.

Barutcu AR, Lajoie BR, Fritz AJ, McCord RP, Nickerson JA, van Wijnen AJ, Lian JB, Stein JL, Dekker J, Stein GS, et al. 2016. SMARCA4 regulates gene expression and higher-order chromatin structure in proliferating mammary epithelial cells. Genome Res gr.201624.115. http://genome.cshlp.org/lookup/doi/10.1101/gr.201624.115.

Botstein D, Fink GR. 2011. Yeast: An experimental organism for 21st century biology. Genetics 189: 695–704.

Bronshtein I, Kepten E, Kanter I, Berezin S, Lindner M, Redwood AB, Mai S, Gonzalo S, Foisner R, Shav-Tal Y, et al. 2015. Loss of lamin A function increases chromatin dynamics in the nuclear interior. Nat Commun 6: 8044. http://www.nature.com/doifinder/10.1038/ncomms9044.

Bystricky K. 2015. Chromosome dynamics and folding in eukaryotes: insights from live cell microscopy. FEBS Lett. http://linkinghub.elsevier.com/retrieve/pii/S0014579315006110.

Cabal GG, Genovesio A, Rodriguez-Navarro S, Zimmer C, Gadal O, Lesne A, Buc H, Feuerbach-Fournier F, Olivo-Marin J-C, Hurt EC, et al. 2006. SAGA interacting factors confine sub-diffusion of transcribed genes to the nuclear envelope. Nature 441: 770–3. http://www.ncbi.nlm.nih.gov/pubmed/16760982 (Accessed August 20, 2014).

Chambeyron S, Bickmore WA. 2004. Chromatin decondensation and nuclear reorganization of the HoxB locus upon induction of transcription. 1119–1130.

Chen B, Gilbert L a, Cimini B a, Schnitzbauer J, Zhang W, Li G-W, Park J, Blackburn EH, Weissman JS, Qi LS, et al. 2013. Dynamic imaging of genomic loci in living human cells by an optimized CRISPR/Cas system. Cell 155: 1479–91. http://www.ncbi.nlm.nih.gov/pubmed/24360272 (Accessed July 9, 2014).

Chuang C-H, Carpenter AE, Fuchsova B, Johnson T, de Lanerolle P, Belmont AS. 2006a. Long-Range Directional Movement of an Interphase Chromosome Site. Curr Biol 16: 825–831.

Chuang CH, Carpenter AE, Fuchsova B, Johnson T, de Lanerolle P, Belmont AS. 2006b. Long-Range Directional Movement of an Interphase Chromosome Site. Curr Biol 16: 825–831.

Cisse II, Izeddin I, Causse SZ, Boudarene L, Senecal A, Muresan L, Dugast-darzacq C, Hajj B. 2013. Polymerase II Clustering in. Science (80-) 245: 664–667.

Dalvai M, Fleury L, Bellucci L, Kocanova S, Bystricky K. 2013. TIP48/Reptin and H2A.Z Requirement for Initiating Chromatin Remodeling in Estrogen-Activated Transcription. PLoS Genet 9.

Darzacq X, Shav-Tal Y, de Turris V, Brody Y, Shenoy SM, Phair RD, Singer RH. 2007. In vivo dynamics of RNA polymerase II transcription. Nat Struct Mol Biol 14: 796–806.

Darzacq X, Yao J, Larson DR, Causse SZ, Bosanac L, de Turris V, Ruda VM, Lionnet T, Zenklusen D, Guglielmi B, et al. 2009. Imaging transcription in living cells. Annu Rev Biophys 38: 173–196.

Feuerborn A, Cook PR. 2015. Why the activity of a gene depends on its neighbors. Trends Genet 31: 483490. http://dx.doi.org/10.1016Zj.tig.2015.07.001.

Fullwood MJ, Liu MH, Pan YF, Liu J, Xu H, Mohamed Y Bin, Orlov YL, Velkov S, Ho A, Mei PH, et al. 2009. An oestrogen-receptor-α-bound human chromatin interactome. Nature 462: 58–64. http://www.nature.com/doifinder/10.1038/nature08497.

Ghamari A, Tte M, Van De Corput PC, Thongjuea S, Van Cappellen WA, Van Ijcken W, Van Haren J, Soler E, Eick D, Lenhard B, et al. 2013. In vivo live imaging of RNA polymerase II transcription factories in primary cells. 767–777.

Giorgetti L, Galupa R, Nora EP, Piolot T, Lam F, Dekker J, Tiana G, Heard E. 2014. Predictive polymer modeling reveals coupled fluctuations in chromosome conformation and transcription. Cell 157: 950–963.

Graham TGW, Wang X, Song D, Etson CM, van Oijen AM, Rudner DZ, Loparo JJ. 2014. ParB spreading requires DNA bridging. Genes Dev 28: 1228–1238.

Gribnau J, de Boer E, Trimborn T, Wijgerde M, Milot E, Grosveld F, Fraser P. 1998. Chromatin interaction mechanism of transcriptional control in vivo. EMBO J 17: 6020–7. http://www.pubmedcentral.nih.gov/articlerender.fcgi?artid=1170928&tool=pmcentrez&rendertype=abstract.

Haddad N, Jost D, Vaillant C. 2017. Perspectives: using polymer modeling to understand the formation and function of nuclear compartments. Chromosom Res. http://link.springer.com/10.1007/s10577-016-9548-2.

Hah N, Danko CG, Core L, Waterfall JJ, Siepel A, Lis JT, Kraus WL. 2011. A rapid, extensive, and transient transcriptional response to estrogen signaling in breast cancer cells. Cell 145: 622–634. http://dx.doi.org/10.1016/j.cell.2011.03.042.

Hajjoul H, Mathon J, Ranchon H, Goiffon I, Alber B, Gadal O, Bystricky K, Bancaud A. 2013. High throughput chromatin motion tracking in living yeast reveals the flexibility of the fiber throughout the genome. Genome Res 23: 1829–1838.

Hancock R. 2004. Internal organisation of the nucleus: assembly of compartments by macromolecular crowding and the nuclear matrix model. Biol Cell 96: 595–601.

Hsieh T-HS, Weiner A, Lajoie B, Dekker J, Friedman N, Rando OJ. 2015. Mapping Nucleosome Resolution Chromosome Folding in Yeast by Micro-C. Cell 162: 108–119. http://linkinghub.elsevier.com/retrieve/pii/S0092867415006388.

Huet S, Lavelle C, Ranchon H, Carrivain P, Victor JM, Bancaud A. 2014. Relevance and limitations of crowding, fractal, and polymer models to describe nuclear architecture. 1st ed. Elsevier Inc. http://dx.doi.org/10.1016/B978-0-12-800046-5.00013-8.

Jonkers I, Kwak H, Lis JT. 2014. Genome-wide dynamics of Pol II elongation and its interplay with promoter proximal pausing, chromatin, and exons. Elife 2014: 1–25.

Kepten E, Weron A, Sikora G, Burnecki K, Garini Y. 2015. Guidelines for the fitting of anomalous diffusion mean square displacement graphs from single particle tracking experiments. PLoS One 10: 1–10. http://dx.doi.org/10.1371/journal.pone.0117722.

Kimura H, Sugaya K, Cook PR. 2002. The transcription cycle of RNA polymerase II in living cells. J Cell Biol 159:777–782.

Kocanova S, Kerr EA, Rafique S, Boyle S, Katz E, Caze-Subra S, Bickmore WA, Bystricky K. 2010. Activation of estrogen-responsive genes does not require their nuclear co-localization. PLoS Genet 6.

Le Dily FL, Ba?? D, Pohl A, Vicent GP, Serra F, Soronellas D, Castellano G, Wright RHG, Ballare C, Filion G, et al. 2014. Distinct structural transitions of chromatin topological domains correlate with coordinated hormone-induced gene regulation. Genes Dev 28: 2151–2162.

Liu X-F, Bagchi MK. 2004. Recruitment of distinct chromatin-modifying complexes by tamoxifen-complexed estrogen receptor at natural target gene promoters in vivo. J Biol Chem 279: 15050–8. http://www.ncbi.nlm.nih.gov/pubmed/14722073 (Accessed October 18, 2014).

Ma H, Naseri A, Reyes-Gutierrez P, Wolfe S a., Zhang S, Pederson T. 2015. Multicolor CRISPR labeling of chromosomal loci in human cells. Proc Natl Acad Sci 201420024. http://www.pnas.org/lookup/doi/10.1073/pnas.1420024112.

Manzo C, Garcia-Parajo MF. 2015. A review of progress in single particle tracking: from methods to biophysical insights. Reports Prog Phys 78: 124601.

Métivier R, Penot G, HÐübner MR, Reid G, Brand H, Koš M, Gannon F. 2003. Estrogen receptor-α directs ordered, cyclical, and combinatorial recruitment of cofactors on a natural target promoter. Cell 115: 751–763.

Mitchell JA, Fraser P. 2008. Transcription factories are nuclear subcompartments that remain in the absence of transcription. Genes Dev 22: 20–25.

Mourad R, Hsu P-Y, Juan L, Shen C, Koneru P, Lin H, Liu Y, Nephew K, Huang TH, Li L. 2014. Estrogen induces global reorganization of chromatin structure in human breast cancer cells. PLoS One 9: e113354. http://www.ncbi.nlm.nih.gov/pubmed/25470140 (Accessed December 5, 2014).

N. Chenouard, I. Bloch JO-M. 2013. Multiple Hypothesis Tracking for Cluttered Biological Image Sequences. IEEE Trans Pattern Anal Mach Intell 35: 2736–3750.

Neumann FR, Dion V, Gehlen LR, Tsai-Pflugfelder M, Schmid R, Taddei A, Gasser SM. 2012. Targeted INO80 enhances subnuclear chromatin movement and ectopic homologous recombination. Genes Dev 26: 369–383.

Ochiai H, Sugawara T, Yamamoto T. 2015. Simultaneous live imaging of the transcription and nuclear position of specific genes. Nucleic Acids Res 1–12. http://nar.oxfordjournals.org/lookup/doi/10.1093/nar/gkv624.

Osborne CS, Chakalova L, Brown KE, Carter D, Horton A, Debrand E, Goyenechea B, Mitchell J a, Lopes S, Reik W, et al. 2004. Active genes dynamically colocalize to shared sites of ongoing transcription. Nat Genet 36: 1065–1071.

P. Thévenaz, U.E. Ruttimann MU. 1998. A Pyramid Approach to Subpixel Registration Based on Intensity. IEEE Trans Image Process 7: 27–41.

Passot FM, Calderon V, Fichant G, Lane D, Pasta F. 2012. Centromere Binding and Evolution of Chromosomal Partition Systems in the Burkholderiales. 194: 3426–3436.

Robinett CC, Straight A, Li G, Willhelm C, Sudlow G, Murray A, Belmont AS. 1996. In vivo localization of DNA sequences and visualization of large-scale chromatin organization using lac operator/repressor recognition. J Cell Biol 135: 1685.

Sanchez A, Cattoni DI, Walter J-C, Rech J, Parmeggiani A, Nollmann M, Bouet J-Y. 2015. Stochastic SelfAssembly of ParB Proteins Builds the Bacterial DNA Segregation Apparatus. Cell Syst 1: 163–173. http://linkinghub.elsevier.com/retrieve/pii/S2405471215000575.

Saxton MJ. 2009. Single Particle Tracking. Fundam Concepts Biophys 1–33.

Sbalzarini IF, Koumoutsakos P. 2005. Feature point tracking and trajectory analysis for video imaging in cell biology. J Struct Biol 151: 182–195.

Schuettengruber B, Cavalli G. 2009. Recruitment of Polycomb group complexes and their role in the dynamic regulation of cell fate choice. Development 136: 3531–3542.

Shang Y, Hu X, Direnzo J, Lazar MA, Brown M, Endocrinology D. 2000. Cofactor Dynamics and Sufficiency in Estrogen Receptor - Regulated Transcription. 103: 843–852.

Stasevich TJ, Hayashi-Takanaka Y, Sato Y, Maehara K, Ohkawa Y, Sakata-Sogawa K, Tokunaga M, Nagase T, Nozaki N, McNally JG, et al. 2014. Regulation of RNA polymerase II activation by histone acetylation in single living cells. Nature 516: 272–275. http://www.nature.com/doifinder/10.1038/nature13714 (Accessed September 21, 2014).

Stavreva D a, Coulon A, Baek S, Sung M, John S, Stixova L, Tesikova M, Hakim O, Miranda T, Hawkins M, et al. 2015. Dynamics of chromatin accessibility and long-range interactions in response to glucocorticoid pulsing. 1–13.

Straight AF, Belmont AS, Robinett CC, Murray AW. 1996. GFP tagging of budding yeast chromosomes reveals that protein-protein interactions can mediate sister chromatid cohesion. Curr Biol 6: 15991608. http://linkinghub.elsevier.com/retrieve/pii/S0960982202707835.

Taddei A, Gasser SM. 2012. Structure and function in the budding yeast nucleus. Genetics 192: 107–129.

Taddei A, Van Houwe G, Hediger F, Kalck V, Cubizolles F, Schober H, Gasser SM. 2006. Nuclear pore association confers optimal expression levels for an inducible yeast gene. Nature 441: 774–8. http://www.ncbi.nlm.nih.gov/pubmed/16760983 (Accessed October 9, 2014).

Therizols P, Illingworth RS, Courilleau C, Boyle S, Wood AJ, Bickmore WA. 2014. Chromatin decondensation is sufficient to alter nuclear organization in embryonic stem cells. 2–7.

Ulianov S V., Khrameeva EE, Gavrilov AA, Flyamer IM, Kos P, Mikhaleva EA, Penin AA, Logacheva MD, Imakaev M V., Chertovich A, et al. 2016. Active chromatin and transcription play a key role in chromosome partitioning into topologically associating domains. Genome Res 26: 70–84.

Vispé S, DeVries L, Créancier L, Besse J, Bréand S, Hobson DJ, Svejstrup JQ, Annereau J-P, Cussac D, Dumontet C, et al. 2009. Triptolide is an inhibitor of RNA polymerase I and II-dependent transcription leading predominantly to down-regulation of short-lived mRNA. Mol Cancer Ther 8: 2780–2790.

Wachsmuth M, Conrad C, Bulkescher J, Koch B, Mahen R, Isokane M, Pepperkok R, Ellenberg J. 2015. High-throughput fluorescence correlation spectroscopy enables analysis of proteome dynamics in living cells. Nat Biotechnol 33: 384–9.

Wang R, Mozziconacci J, Bancaud A, Gadal O. 2015. Principles of chromatin organization in yeast: relevance of polymer models to describe nuclear organization and dynamics. Curr Opin Cell Biol 34: 54–60. http://linkinghub.elsevier.com/retrieve/pii/S0955067415000447.

Yunger S, Rosenfeld L, Garini Y, Shav-Tal Y. 2010. Single-allele analysis of transcription kinetics in living mammalian cells. Nat Methods 7: 631–633. http://dx.doi.org/10.1038/nmeth.1482.

